# Identification of Animal Behavioral Strategies by Inverse Reinforcement Learning

**DOI:** 10.1101/129007

**Authors:** Shoichiro Yamaguchi, Honda Naoki, Muneki Ikeda, Yuki Tsukada, Shunji Nakano, Ikue Mori, Shin Ishii

**Affiliations:** Graduate School of Informatics, Kyoto University, Sakyo, Kyoto, Kyoto, Japan; Graduate School of Biostudies, Kyoto University, Sakyo, Kyoto, Kyoto, Japan; Research Center for Dynamic Living Systems, Kyoto University, Sakyo, Kyoto, Kyoto, Japan; Graduate School of Science, Nagoya University, Chikusa, Nagoya, Aichi, Japan

**Keywords:** Behavioral Strategy, Reinforcement learning, Reward, Machine learning, Optimal control, LMDP, KL control, C. elegans, Thermotaxis

## Abstract

Animals are able to reach a desired state in an environment by controlling various behavioral patterns. Identification of the behavioral strategy used for this control is important for understanding animals’ decision-making and is fundamental to dissect information processing done by the nervous system. However, methods for quantifying such behavioral strategies have not been fully established. In this study, we developed an inverse reinforcement-learning (IRL) framework to identify an animal’s behavioral strategy from behavioral time-series data. As a particular target, we applied this framework to *C. elegans* thermotactic behavior; after cultivation at a constant temperature with or without food, the fed and starved worms prefer and avoid from the cultivation temperature on a thermal gradient, respectively. Our IRL approach revealed that the fed worms used both absolute and temporal derivative of temperature and that their strategy comprised mixture of two strategies: directed migration (DM) and isothermal migration (IM). The DM is a strategy that the worms efficiently reach to specific temperature, which explained thermotactic behaviors of the fed worms. The IM is a strategy that the worms track along a constant temperature, which reflects isothermal tracking well observed in previous studies. We also showed the neural basis underlying the strategies, by applying our method to thermosensory neuron-deficient worms. In contrast to fed animals, the strategy of starved animals indicated that they escaped the cultivation temperature using only absolute, but not temporal derivative of temperature. Thus, our IRL-based approach is capable of identifying animal strategies from behavioral time-series data and will be applicable to wide range of behavioral studies, including decision-making of other organisms.

**Author Summary:** Understanding animal decision-making has been a fundamental problem in neuroscience and behavioral ecology. Many studies analyze actions that represent decision-making in behavioral tasks, in which rewards are artificially designed with specific objectives. However, it is impossible to extend this artificially designed experiment to a natural environment, because in a natural environment, the rewards for freely-behaving animals cannot be clearly defined. To this end, we must reverse the current paradigm so that rewards are identified from behavioral data. Here, we propose a new reverse-engineering approach (inverse reinforcement learning) that can estimate a behavioral strategy from time-series data of freely-behaving animals. By applying this technique with thermotaxis in *C. elegans*, we successfully identified the reward-based behavioral strategy.

## Introduction

Animals develop behavioral strategies, a set of sequential decisions necessary for organizing appropriate actions in response to environmental stimuli, to ensure their survival and reproduction. Such strategies lead the animal to its preferred states and provide them with effective solution to overcome difficulties in a given environment. For example, foraging animals are known to optimize their strategy to most efficiently exploit food sources [1]. Therefore, understanding behavioral strategies of biological organisms is important from biological, ethological, and engineering point of views.

A number of behavioral studies have recorded behavioral sequences that should reflect the animal’s overall behavioral strategies. However, mechanistic description such as behavioral strategy is on a different level of understanding from phenomenological description of the recorded behavior [2], and a method that can objectively identify behavioral strategy, a mechanistic component of the behavior, based on behavioral time-series data has not been well established. From a theoretical viewpoint, this mechanistic component corresponds to an algorithmic/representational level of understanding for information processing system [3]. To derive behavioral strategy from quantitative time-series of behavioral data, we propose a new computational framework based on the concept of reinforcement learning (RL).

RL is a mathematical paradigm to represent how animals adaptively learn behavioral strategies to maximize cumulative rewards via trial and error [4] (blue arrow in Fig 1A). Previous study indicated that dopamine neural activity represents prediction error of reward [5], similar to temporal difference (TD) learning in RL [6], suggesting that RL-based regulation underlies animal’s behavioral learning. Even in a simple neural circuit in *Caenorhabditis elegans*, dopamine-dependent neural circuit involved in the exploring behavior is reminiscent of RL [7]. Thus some animal’s behavioral strategies are likely to be associated with reward in its neural system.

**Fig 1:**
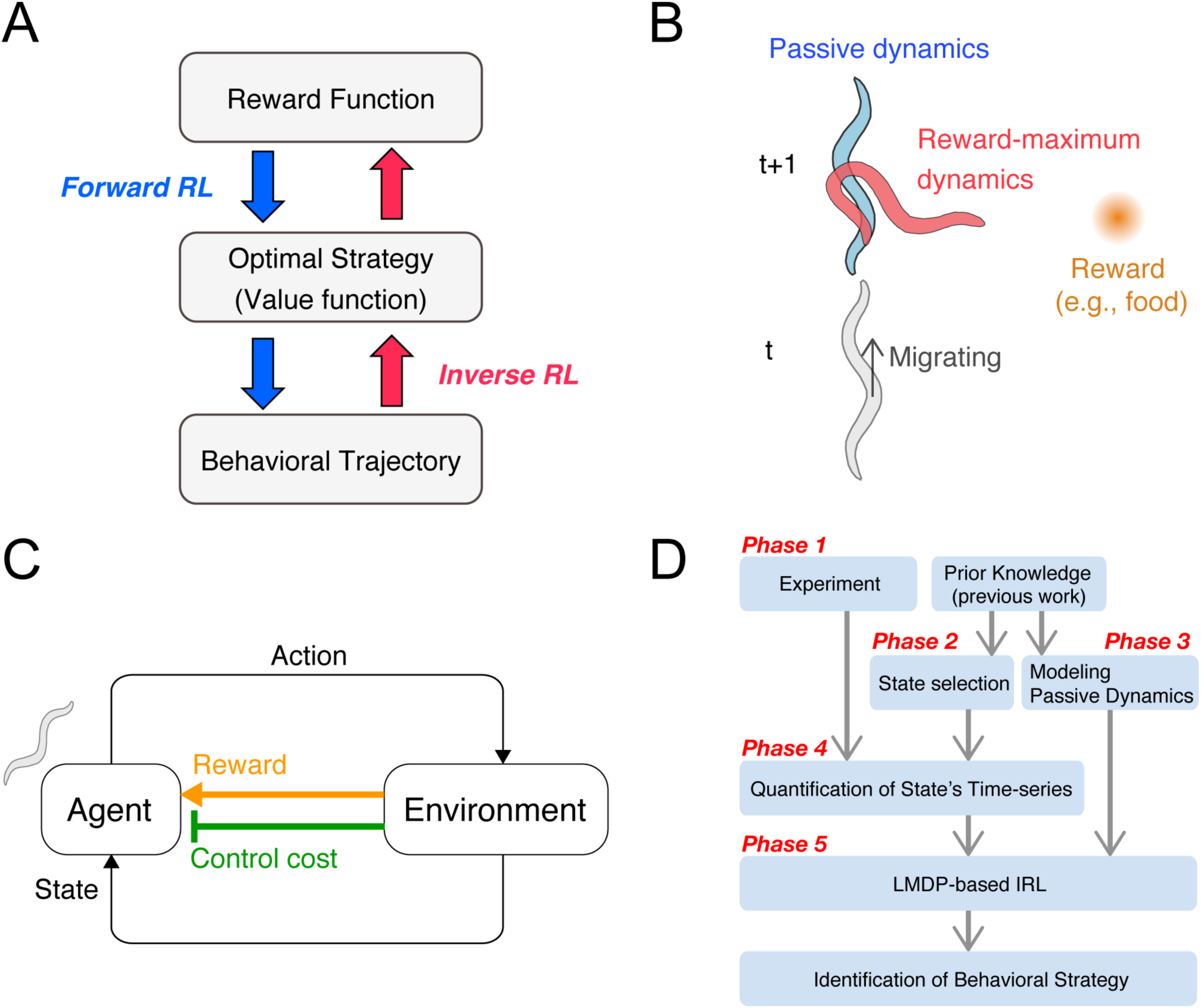
Concept and procedure of the inverse reinforcement learning (IRL)-based approach. **(A)** RL represents a forward problem, in which a behavioral strategy is determined to maximize the cumulative reward given as a series of state-dependent rewards. IRL represents an inverse problem, in which a behavioral strategy, or its underlying value and reward functions, is estimated in order to reproduce an observed series of behaviors. The behavioral strategy is evaluated by the profiles of the identified functions. **(B)** Examples of passive dynamics and controlled dynamics. Here, an animal migrates upwards whereas the food (reward) is placed to its right. In this situation, if the animal continues to migrate upwards, the distance to the food increases. If the animal exercises harder body control, that is, changes its migrating direction toward the food, the distance to the food decreases. The animal should therefore make decisions based on a tradeoff between these two dynamics. **(C)** The agent-environment interaction. The agent autonomously acts in the environment, observes the resultant state-transition through its sensory system, and receives not only the state reward but also the body control cost. The behavioral strategy is optimized in order to maximize the accumulation of the net reward, which is given as state reward minus body control cost. **(D)** Guideline for the IRL framework. This guideline outlines the general procedures of the IRL framework for the identification of animal behavioral strategies. Details are explained in the main text.

Inverse reinforcement learning (IRL) is a recently-developed machine learning framework that can solve an inverse problem of RL (blue arrow in **Fig 1A**) and estimate reward-based strategy from behavioral time-series [8,9]. One engineering application of IRL is apprenticeship learning. For example, the seminal studies on IRL employed a radio-controlled helicopter, for which the state-dependent rewards of an expert were estimated by using the observed time-series of both the human expert’s manipulation and the helicopter’s state. Consequently, autonomous control of the helicopter was achieved by (forward) RL that utilized the estimated rewards [10,11]. This engineering application prompted studies in which IRL was used to identify the behavioral strategies of animals. Recently, application studies of IRL have emerged, mostly regarding human behavior, with a particular interest in constructing artificially intelligent systems that mimic human behavior [12–15]. In these studies, the experimenters designed the behavioral tasks with specific objectives, and the observed behavioral strategies are therefore usually expected. However, IRL applications to freely behaving animals in a more natural environment are far from established.

In an effort to apply the IRL to freely behaving animals, we chose thermotaxis of *C. elegans* as a model behavior regulated by specific strategies. If worms are cultivated at a constant temperature with plenty of food and placed on a thermal gradient without food, the fed worms show appetitive response to the cultivation temperature [16,17]. On the other hand, if worms are cultivated at a constant temperature without food, the starved worms aversively avoid the cultivation temperature on a thermal gradient [18,19]. Despite that the worms are not aware of neither the spatial temperature profile nor their current location, it is obvious that worms somehow make rational decisions, depending on fed or starved conditions. Although there are multiple candidates of strategies that can theoretically lead animals to the goals, behavioral strategies that animals utilize in each condition are largely unknown, because stochastic nature of behavioral sequences conceals a principle of behavioral regulation, as in the case of many other animal’s behaviors.

In this study, we developed a new IRL framework to identify the behavioral strategy as a value function. The value function represents benefit of each state, namely, how much future rewards are expected starting from the state. Applying this IRL framework to behavioral time-series data of freely migrating *C. elegans*, we identified the value functions underlying the thermotactic strategies. The strategy of fed animals was based on sensory information of both absolute and temporal derivative of temperature, and comprised mixture of two modes, which correspond to directed migration to the cultivation temperature and isothermal migration along contour at constant temperature, respectively. The starved worm, on the other hand, used only absolute temperature but not its temporal derivative for escaping the cultivation temperature. By further applying the IRL to thermosensory neuron-impaired worms, we found that the AFD neuron is fundamental to the directed migration of the fed strategy. Thus, our IRL framework can reveal what is most preferable, optimal state for animals and, more importantly, how animals reach the state, thereby providing clues for understanding the computational principles in the nervous system.

## Results

### Heart of the IRL framework

To identify an animal’s behavioral strategy based on IRL, we initially make the assumption that the animal’s behaviors are the result of a balance between two factors: passive dynamics (blue worm in **Fig 1B**) and reward-maximizing dynamics (red worm in **Fig 1B**). These factors correspond to inertia-based and purpose-driven body movements, respectively. Even if a worm moving in a straight line wants to make a purpose-driven turn towards a reward, it cannot turn suddenly, due to the inertia of its already moving body. Thus, it is reasonable to consider that an animal’s behaviors are optimized by taking both factors into account, namely by minimizing resistance to the passive dynamics and maximizing approach to the destination (reward). Such a behavioral strategy has recently been modeled as a linearly-solvable Markov decision process (LMDP) [20], in which the agent requires not only a state-dependent reward, but also a control cost for quantifying resistance to the passive dynamics (**Fig 1C**) (see **Materials and Methods**). Importantly, the optimal strategy in the LMDP is analytically obtained as a probability of controlled state transition [20]:

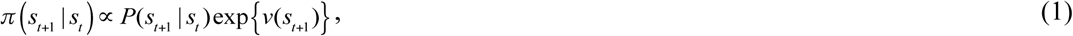

where *s*_*t*_ and *v*(*s*) indicate the animal’s state at time step *t* and a value function defined as the expected sum of state-dependent rewards, *r*(*s*), and negative control cost, *KL*[*π* (· | *s*) ‖ *p*(· | *s*)], from state *s* toward the future, respectively; *P*(*s*_*t+*1_|*s*_*t*_) represents a probability of uncontrolled state transition, which represents the passive dynamics from *s*_*t*_ to *s*_*t+*1_. In this equation, the entire set of *v*(*s*) represents the behavioral strategy. Thus, the identification of the animal’s behavioral strategy is equivalent to an estimation of the value function *v*(*s*) based on observed behavioral data (*s*_*1*_, *s*_*2*_,…*s*_*t*_,…*s*_*T*_) (red arrow in **Fig 1A**). This estimation was performed by maximum likelihood estimation (MLE) [21], and is an instance of IRL. Note that in this estimation, we introduced a constraint to make the value function smooth, because animals generate similar actions in similar states. This constraint is essential to stably estimate the behavioral strategy of animals (see **Materials and Methods**). In summary, identification of an animal’s behavioral strategy from behavioral data requires only the appropriate design of state representation and passive dynamics.

### Guidelines for the application of the IRL framework

The guideline of the IRL was depicted in flowchart (**Fig 2D**). Suppose one is interested in a certain animal’s behavioral strategy. The first step would be to perform a behavioral experiment (**phase 1 in Fig 1D**), which can be either a freely-moving task or a conditional task. The second step would then be to select the states, *s*, and model the passive dynamics, *P*(*s*_*t+*1_|*s*_*t*_), based on which the animal develops its strategy (**phase 2 and 3 in Fig 1D**). At this time, prior knowledge about the kinds of sensory information the animal processes provides useful information for the appropriate selection of the states and the passive dynamics. The third step is the quantification of the time-series of selected states (**phase 4 in Fig 1D**). The fourth step is the implementation of the LMDP-based IRL to estimate the value function (**phase 5 in Fig 1D**). At this point, the behavioral strategy can be identified. Following this guideline, we applied the IRL framework to freely-migrating *C. elegans* under a thermal gradient.

**Fig 2:**
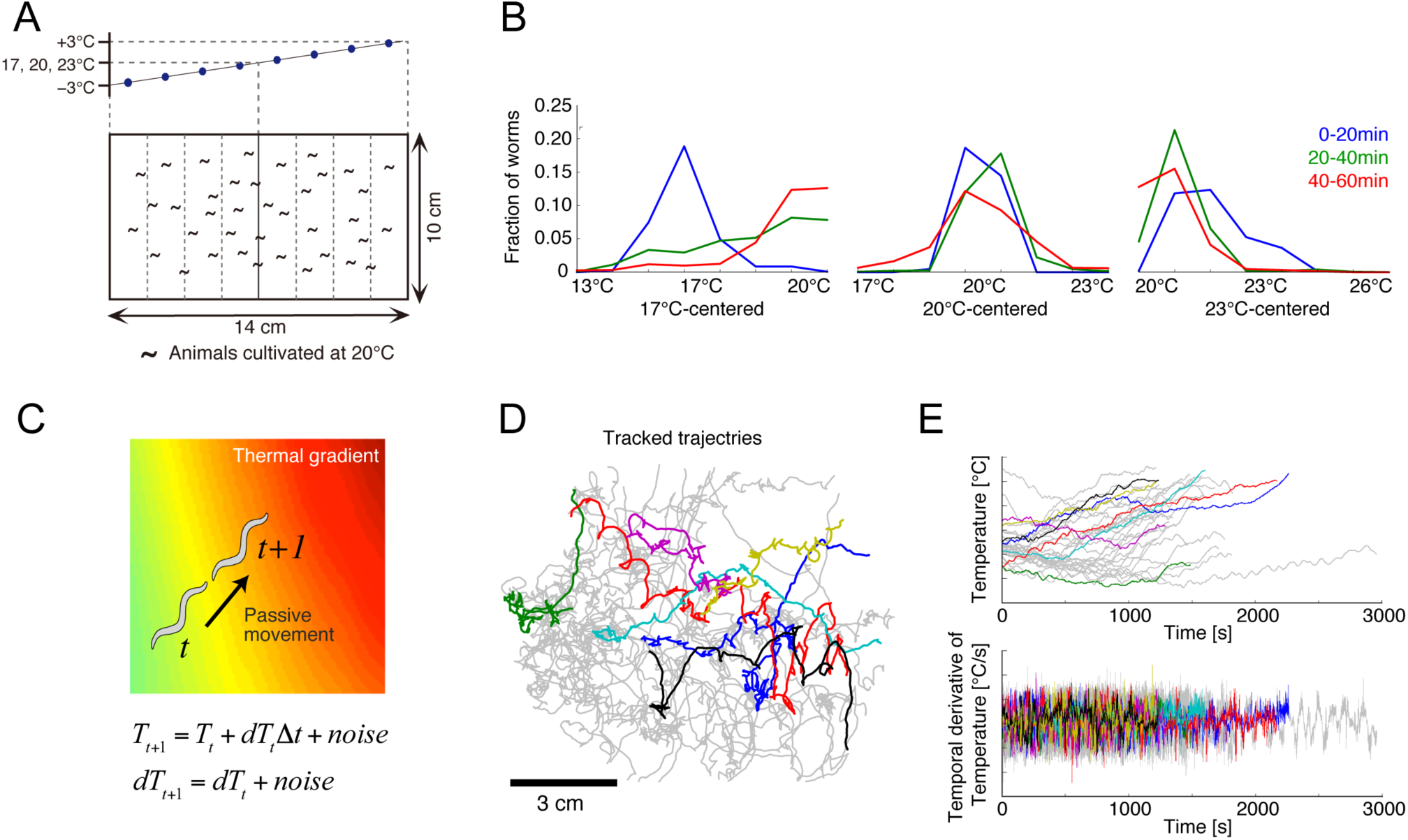
Thermotactic behavior of *C. elegans*. **(A)** Thermotaxis assays with a thermal gradient. In each assay, a linear temperature gradient was set along the agar surface, whose center was set at either 17, 20, or 23 °C. At the onset of the assays, fed or starved worms were uniformly placed on the agar surface. **(B)** Representative trajectories of worms extracted by the multi-worm tracking system (n=33 in this panel). Different colors indicate different individual worms. **(C)** Passive dynamics of persistent migration on a linear thermal gradient. **(D)** Temporal changes in the worms’ spatial distribution under the 17 °C-, 20 °C- and 23°C-centered thermal gradients in the fed condition. **(E)** Time series of the temperature and its derivative experienced by the migrating worms shown in C (colors correspond to those in D).

### *Phase 1: Experiment on* C. elegans *thermotaxis*

To identify the behavioral strategy underlying the thermotactic behavior of *C. elegans*, we performed population thermotaxis assays, in which 80–150 worms that had been cultivated at 20 °C were placed on the surface of an agar plate with controlled thermal gradients (**Fig 2A**). Behavioral crosstalk is negligible, because the rate of physical contact is low at this worm density. We prepared three different thermal gradients centered at 17 °C, 20 °C, and 23 °C, to collect behavioral data; these gradients would encourage ascent up the gradient, movement around the center, and descent down the gradient, respectively. We confirmed that the fed worms aggregated around the cultivation temperature in all gradients (**Fig 2B**).

### *Phase 2: Selection of* C. elegans’ *states*

We here defined state, which is signified by *s* in equation (1). The state must represent the sensory information that the worms process during thermotaxis. Because previous studies have shown that the thermosensory neuron AFD encodes the temporal derivative of temperature [22,23], we assumed that the worm made decisions in order to select appropriate actions (i.e., migration direction and speed) based not only on temperature, but also on its temporal derivative. We then represented a state by a two-dimensional (2D) sensory space, *s*=(*T, dT*), where *T* and *dT* denote temperature and its temporal derivative, respectively. This means that the value function in equation (1) is given as a function of *T* and *dT, v*(*T, dT*). Note that we did not select spatial coordinates on the assay plate as the state, because the worms are not able to recognize neither the spatial temperature profile nor their current position on the assay plate.

### *Phase 3: Modeling* C. elegans’ *passive dynamics*

We also defined passive dynamics, which is signified by *P*(*s*_*t+*1_|*s*_*t*_) in equation (1). The passive dynamics are given by state transitions as a consequence of uncontrolled behavior. We assumed that a worm likely migrates in a persistent direction, but in a sometimes fluctuating manner. During state transition in a short time interval, the thermal gradient in local area can be addressed as linear (**Fig. 2C**). Thus, it is reasonable to model the passive transition from a state *s*_*t*_=(*T*_*t*_, *dT*_*t*_) to the next state *s*_*t+*1_=(*T*_*t+*1_, *dT*_*t+*1_), where *dT*_*t+*1_ maintains *dT*_*t*_ with white noise and *T*_*t+*1_ is updated as *T*_*t*_*+dT*_*t*_ with white noise. Accordingly, *P*(*s*_*t+*1_|*s*_*t*_) is to be simply expressed by a normal distribution (see **Materials and Methods**). Please note the distinction between *T* and *t* throughout this paper.

### Phase 4: Quantification of time-series of thermosensory states

To quantify thermosensory states selected in phase 2, we tracked the trajectories of individual worms over 60 min within each gradient, using multi-worm tracking software [24] (**Fig 2D**). We then obtained time-series of the temperature that each individual worm experienced (upper panel in **Fig 2E**). We also calculated the temporal derivative of the temperature using a Savitzky-Golay filter [25] (lower panel in **Fig 2E**). State trajectories in the *T*-*dT* space were also plotted (**S2A Fig**).

### Phase 5: Thermotactic strategy identified by IRL

Using the collected time-series of state, *s*=(*T, dT*), and passive dynamics, *P*(*s*_*t+*1_|*s*_*t*_), we performed IRL (the estimation of the value function *v*(*s*)). We modified an existing estimation method called OptV [21] by introducing a smoothness constraint (see **Materials and Methods**), and confirmed that this smoothness constraint was indeed effective in accurately estimating the value function when applied to artificial data simulated by equation (1) (**S1 Fig**). Since the method was able to powerfully estimate a behavioral strategy based on artificial data, we next applied it to the behavioral data of fed *C. elegans*.

Our method successfully estimated the value function of *T* and *dT, v*(*T, dT*) (**Fig 3A**), and visualized exp(*v*(*T, dT*)), called the desirability function [21] (**Fig 3B**). The reward function could be calculated from the identified desirability function using equation (8) (**Fig 3C**). The reward function primarily represents the worms’ preference; the desirability function represents the behavioral strategy, and is thus a result of optimizing cumulative of rewards and negative control costs. Taken together, our method quantitatively clarified the behavior strategy of fed *C. elegans.*

**Fig 3:**
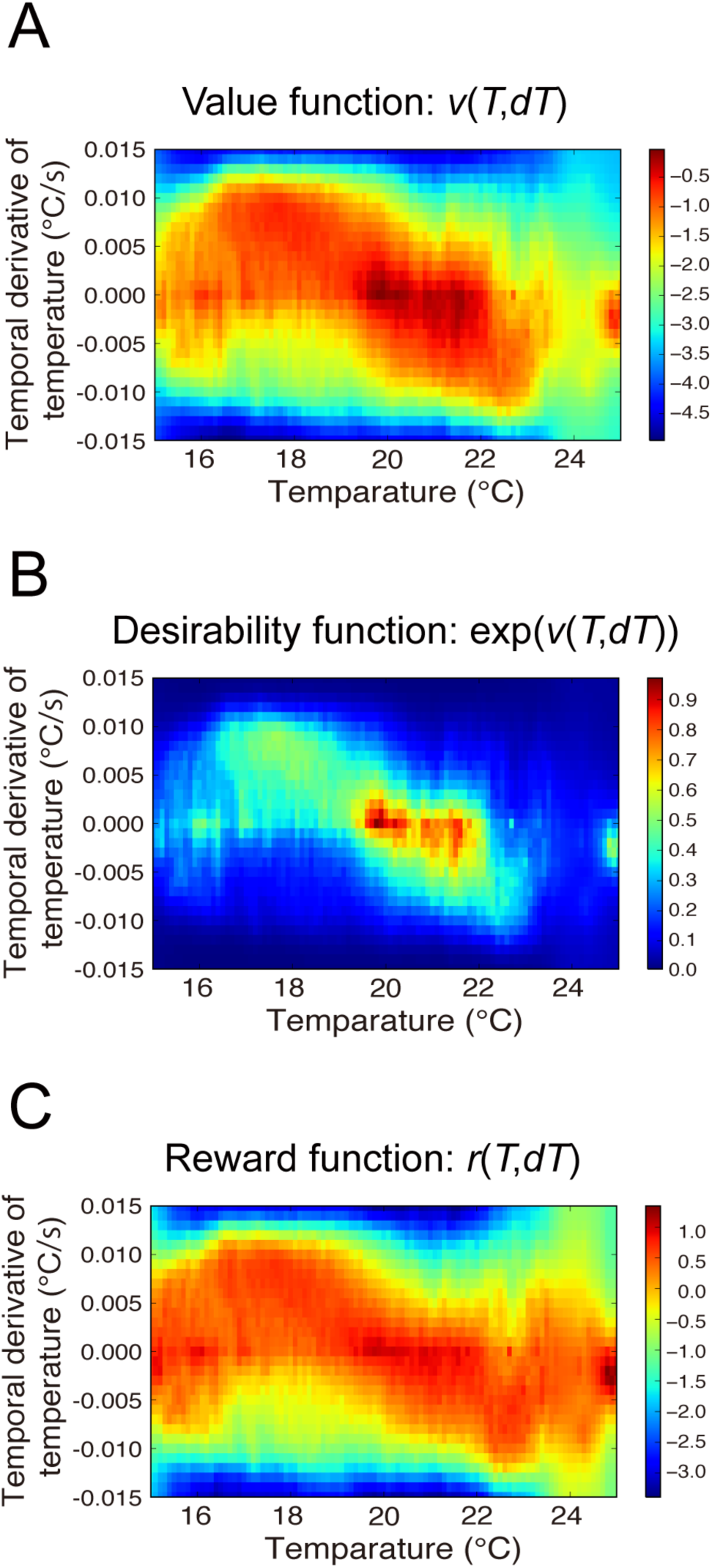
Behavioral strategy identified for fed WT worms. The behavioral strategies of fed WT worms represented by the value **(A)**, desirability **(B)**, and reward **(C)** functions. The worms prefer and avoid the red- and blue-colored states, respectively.

### Interpretation of the identified strategy

Because both the value and desirability functions essentially represent the same thermotactic strategy, we discuss only the desirability function below. We found that the identified desirability function peaked at *T*=20 (°C) and *dT*=0 (°C/s), encouraging the worms to reach and stay close to the cultivation temperature; moreover, we recognized both diagonal and horizontal components (**Fig 3B**), though the latter one was partially truncated by data limitation and data inhomogeneity (**S2B Fig**). The diagonal component represents directed migration (DM), a strategy that enables the worms to efficiently reach the cultivation temperature. For example, at lower temperatures than the cultivation temperature a more positive *dT* is favored, whereas at higher temperature a more negative *dT* is favored. This DM strategy is consistent with the observation that the worms migrate toward the cultivation temperature, and also clarifies how the worms control their thermosensory state throughout migration, which has not been known until now (see **Fig 5** and Discussion). The horizontal component represents isothermal migration (IM), which explains a well-known characteristic called isothermal tracking; the worms typically exhibit circular migration under a concentric thermal gradient [17]. Note that although we used a linear, not a concentric gradient in our thermotaxis assay, our IRL algorithm successfully extracted the isothermal tracking-related IM strategy. We further revealed that the IM strategy worked not only at the cultivation temperature, but also at other temperatures. It must be stressed that the desirability function (**Fig 3B**) controls the strategy of state transition (equation (1)), while the state distribution of *T* and *dT* (**S2B Fig**) was an outcome of the strategy, so that the desirability function was not equivalent to the actual state distribution.

**Fig 4:**
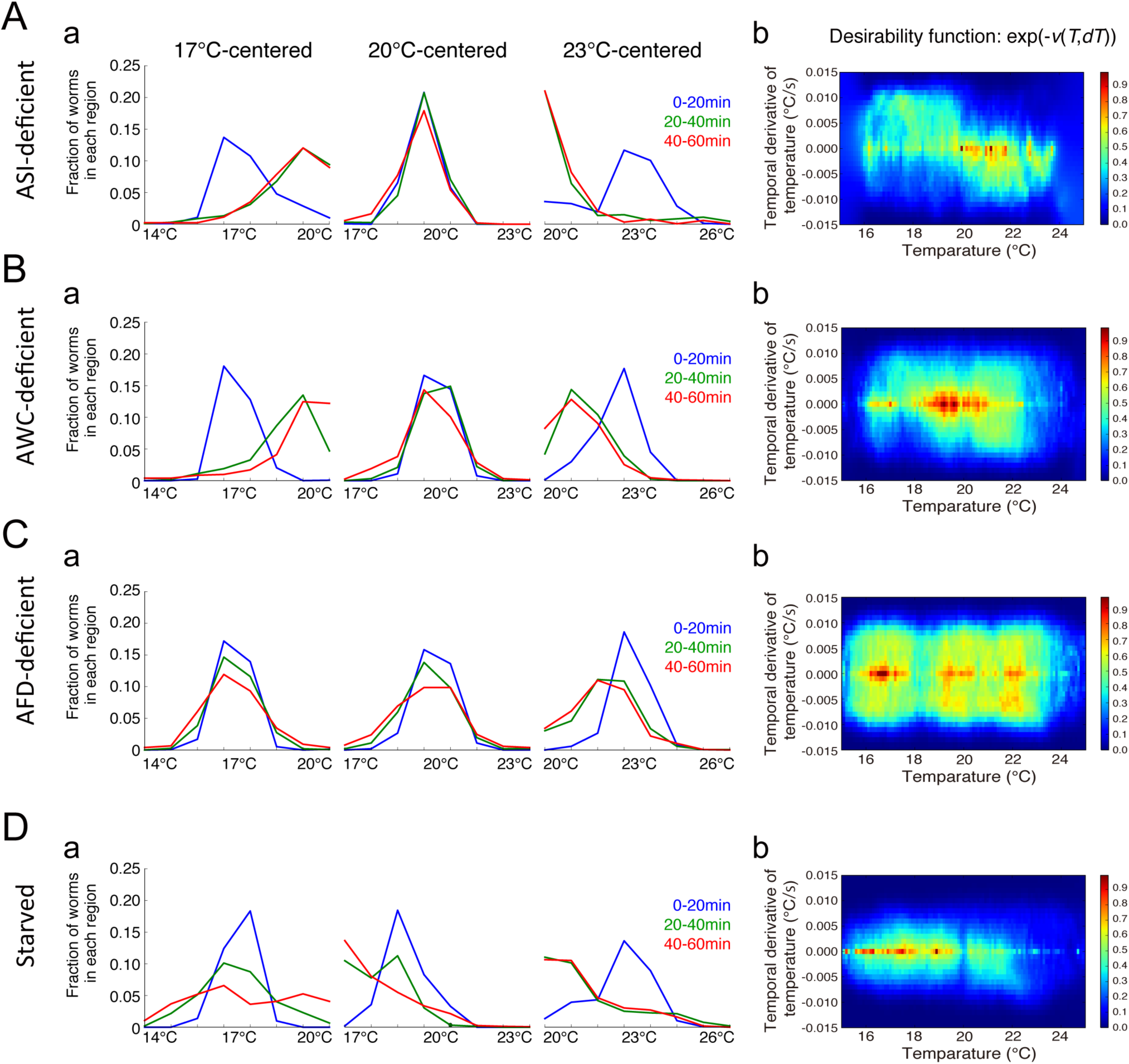
Inverse reinforcement learning (IRL) analyses of ASI-, AWC-, and AFD-deficient worms and starved worms. Temporal changes in distributions of ASI-deficient worms, AWC-deficient worms, AFD-deficient worms and starved worms in the 17 °C-, 20 °C- and 23 °C-centered thermal gradients after the behavior onset are presented in column **a** of A, B, C, and D, respectively. Starved worms disperse under a thermal gradient; AWC-deficient worms migrate to the cultivation temperature, similarly to fed WT worms, and AFD-deficient worms show cryophilic thermotaxis. Corresponding desirability functions are shown in column **b** of A, B, C, and D.

**Fig 5:**
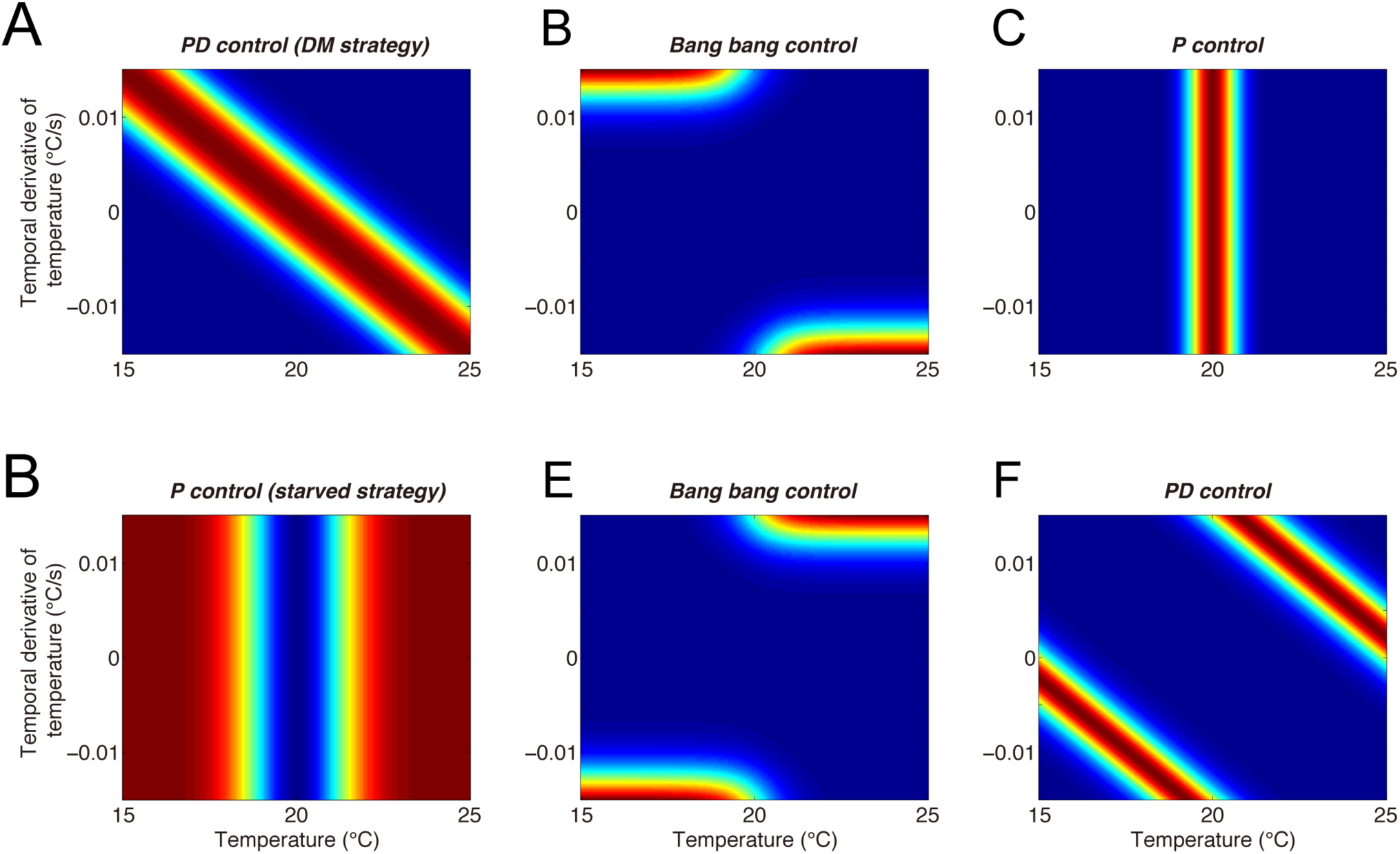
Possible strategies of preferring and avoiding the cultivation temperature. Each panel represents the desirability function of a possible strategy. The prior knowledge that fed worms navigate to the cultivation temperature and starved worms escape the cultivation temperature allowed to propose several possible strategies, but not to identify the worms’ actual strategy (fed worms: **A-C**, starved worms: **E-G**). Our approach identified that the fed worms used the proportional-derivative (PD) control-like DM strategy shown in in), while the starved worms used the proportional (P) control-like strategy shown in (D).

During thermotaxis, the worms basically alternate ‘runs’ and ‘sharp turns’, which correspond to persistent migration with slightly changing direction during long interval and intermittent directional change with large angle during short interval, respectively [26]. Because number of data points during the runs is much larger than those during the sharp turns in total, our IRL could recapitulate a strategy of shallow turns, but not of sharp turns. Indeed, we could not find relationship between the desirability function and rates of sharp turns (**S2C and S2D Fig**).

### Reliability of the identified strategy

We verified the reliability of the identified strategies by following four ways. First, we examined the dimension of the strategy. We estimated the behavioral strategy based on a one-dimensional (1D) state representation, i.e., *s*=(*T*). Comparing 1D and 2D cases, we used cross-validation (see **Materials and Methods**) to confirm that the prediction ability for a future state transition in the 2D behavioral strategy was significantly higher than that in the 1D behavioral strategy (*p=*0.0002; Mann-Whitney U test) (**S3 Fig**). This result indicates that the fed worms utilize sensory information of both absolute and temporal derivative of temperature for their behavioral strategy.

Second, we confirmed whether our IRL approach recapitulated nature of thermotactic behaviors. We simulated temperature trajectories started from 15°C, 20°C and 25°C by sampling state transition from equation (1) using the identified value function. The simulated worm population converged around the cultivation temperature (**S4 Fig**), showing that the identified strategy indeed represents thermotactic property of the fed worms.

Third, we statistically tested the identified DM and IM strategies. As a null hypothesis, we assumed that the worms randomly migrated under a thermal gradient with no behavioral strategy. By means of surrogate method-based statistical testing (see **Materials and Method**), we showed that the DM and IM strategies could not be obtained by chance, indicating that both strategies reflected an actual strategy of thermotaxis (**S5 Fig**).

Fourth, we double checked the DM and IM strategies by performing IRL on another *C. elegans* strain. To this end, we used worms in which chemosensory neuron ASI is genetically ablated via cell-specific expression of caspases [27]. This ASI-deficient appeared to show normal thermotaxis (**Fig. 4Aa**), suggesting that the ASI was not responsible for thermotaxis in our assay. We found clear diagonal and horizontal components in the desirability function, which supports the existence of the DM and IM strategies (**Fig. 4Ab**).

### Strategies of thermosensory neuron-deficient worms

Next, to examine roles of thermosensory system in the strategy, we created two strains in which one of the two thermosensory neurons, AWC or AFD, were deficient [16,17,28]. AWC and AFD had been genetically ablated via cell-specific expression of caspases (see **Materials and Methods**). The AWC-deficient worms appeared to show normal thermotaxis (**Fig 4Ba**). We obtained the desirability functions similar to that of wild type (WT) animals (**Fig 4Bb**), suggesting that the AWC did not play an essential role in thermotaxis.

The AFD-deficient worms in contrast demonstrated cryophilic thermotaxis (**Fig 4Ca**). The desirability function consistently increased with a decrease in temperature (**Fig 4Cb**). We further found that the desirability function lacked the *dT-*dependent structure, indicating that the DM strategy observed in the WT worms had disappeared. Moreover, the fact that the AFD encodes temporal derivatives of temperature [22,23] further corroborates the loss of the *dT-*dependent structure. These results provide an interesting suggestion that the AFD-deficient worms inefficiently aimed for lower temperatures by a strategy primarily depending on the absolute temperature *T*, but not on the temporal derivative of the temperature, *dT* (**Fig 4Cb**). Taken together, our IRL approach clarified that the AFD, but not the AWC neuron, is essential for efficiently navigating to the cultivation temperature.

### Strategy of starved worms

Moreover, we performed IRL on behavioral data from starved worms, which were cultivated at 20°C without food. The starved worms dispersed in low-temperature region and avoided high-temperature region (**Fig 4Da**). In the desirability function, we found the diagonal structure was abandoned (**Fig 4Db**), compared with the desirability function of the fed WT worms (**Fig 3B**), suggesting that the starved worms were not using directed migration. In addition, we noticed that the IM strategy could still be observed (**Fig 5Ab**), which is the first finding that the starved worms retain the ability to perform isothermal tracking. Most importantly, the desirability function at the cultivation temperature was lower than at surrounding temperatures, showing that unlike the fed worm, the starved worms escaped the cultivation temperature region based on sensory information of only the absolute temperature, but not of the temporal derivative of temperature. These results indicate that our method could distinguish between strategies of normally fed and starved *C. elegans*.

## Discussion

We proposed an IRL framework to identify animals’ behavioral strategies based on collected behavioral time-series data. We validated the framework using artificial data, and then applied it to *C. elegans* behavioral data collected during thermotaxis experiments. We quantitatively identified the thermotactic strategies, and discovered that the fed worms used both absolute and temporal derivative of temperature, whereas the starved worms used only absolute temperature. We then visualized the properties of the thermotactic strategy in terms of the desirability function, which successfully identified what states are pleasant and unpleasant for *C. elegans*. Finally, we demonstrated the ability of this technique to discriminate alterations in components within a strategy by comparing the desirability functions of two strains of the worm with impaired thermosensory neuron function; we found that the AFD neuron (but not the AWC) is fundamental to efficiently guided navigation to the cultivation temperature.

### Advantages of our IRL approach

Our IRL approach has three advantages as follows. First, it is generally applicable to behavioral data not only of *C. elegans*, but to that of any animal, by following our guideline (**Fig 1D**). Second, this approach can be applied independently of experimental conditions. Our approach is especially suitable for analyzing behavior in natural conditions where target animals are behaving freely. To the best of our knowledge, this is the first study to identify the behavioral strategy of a freely-behaving animal by IRL. Third, this approach estimates the strategy generating natural behaviors by introducing the passive dynamics in the LMDP. Animals’ movements are usually restricted by external constraints such as inertia and gravity, and by internal (musculoskeletal) constraints, so that animals prefer entering natural, unrestricted state transition. Thus, the LMDP-based IRL is suited for modeling the animals’ behavioral strategy. Although there are several studies of IRL application to human behaviors [12–15], none of these takes account for the passive dynamics. In an era when high-throughput experiments produce massive amounts of the behavioral data, our IRL approach has the potential to become a fundamental tool with applicability in broad behavioral science, including ecology and ethology.

### Validity of the identified strategies

We applied our IRL approach to several cases of worms (WT and three deficient strains), and confirmed that the identified behavioral strategies, that is, the desirability functions, showed no discrepancy in thermotactic behaviors. Fed WT worms aggregated at the cultivation temperature (**Fig 2B**), which can be explained by highest amplitudes of the desirability function at the cultivation temperature (**Fig 3B**). Starved WT worms dispersed around the cultivation temperature (**Fig 4Da**), accompanied by lowered amplitudes of the desirability function at the cultivation temperature (**Fig 4Db**). The ASI- and AWC-deficient worms showed normal thermotaxis (**Fig 4Aa and 4Ba**), and their desirability functions (**Fig 4Ab and 4Bb**) were similar to that of WT animals (**Fig 3B**). The AFD-deficient worms demonstrated cryophilic thermotaxis (**Fig 4Ca**), consistent with higher amplitudes of the desirability function at lower temperatures (**Fig 4Cb**). In summary, these results demonstrate the validity of our approach as well as the potential of the method to determine changes in behavioral strategy.

### Findings beyond what is already known

The identification of the strategies by our IRL approach provides a novel insight into how the *C. elegans* reaches the target temperature on a thermal gradient. We emphasize that in theory, the identified strategy was not the sole solution that was possible for animals to take in order to reach the target state, but there are considered several alternative solutions that would allow animals to navigate to their behavioral goals. Here, we discuss fed and starved strategies while listing possible alternative candidates in terms of control theory [29].

First, regarding the DM strategy of the fed worms (**Fig 5A**), we could provide several examples of alternative strategies that also enable the worm to navigate to the goal (cultivation) temperature. **Fig 5B** shows the desirability function for the worms switching their preference between a positive and a negative gradient in temperatures lower and higher than the goal temperature, namely “bang-bang control”. A previous computational study modeled *C. elegans* thermotaxis based on the bang-bang control [30], in which straight runs and random turnings (i.e., corresponding to omega and reversal turns) were alternated and the run duration was regulated by the temperature, its temporal derivative and the cultivation temperature. **Fig 5C** shows the resulting desirability function if the worms simply prefer the goal temperature, regardless of the temporal derivative of the temperature. This might seem like “proportional (P) control”. However, the identified DM strategy is based on both the absolute temperature and its temporal derivative as shown in **Fig 5A**, so that the worms in fact perform “proportional-derivative (PD) control”, which should be more sophisticated than the bang-bang control.

Second, regarding the strategy of the starved worms (see **Fig 5D**), as discussed above, there are several possible alternative strategies. The worms could escape the cultivation temperature by performing “bang-bang control” or “PD control”, as shown in **Fig 5E and F**. The identified starved strategy is however closer to “P control”, which only uses the absolute temperature. Our IRL-based approach is therefore able to clarify how the worms control their thermosensory state throughout migration, which has not been understood until now.

### Functional significance of DM and IM strategies

We found that the WT worms used a thermotactic strategy consisting of two components, DM and IM strategies (**Fig 3B and Fig 4Ab**). What is the functional meaning of these two strategies? We propose that the existence of these two strategies could be interpreted in terms of balancing exploration and exploitation. Exploitation is the use of pre-acquired knowledge in an effort to obtain rewards, and exploration is the effort of searching for possibly greater rewards. For example, the worm knows that food is associated with the cultivation temperature, and it can exploit that association. On the other hand, the worm could explore different temperatures to search for a larger amount of food than what is available at the cultivation temperature. In an uncertain environment, animals usually face an “exploration-exploitation dilemma” [31]; exploitative behaviors reduce the chance to explore for greater rewards, whereas exploratory behaviors disrupt the collection of the already-available reward. Therefore, an appropriate balance between exploration and exploitation is important for controlling behavioral strategies. We propose the hypothesis that the DM strategy generates exploitative behaviors, whereas the IM strategy generates explorative ones: the worms, through the DM strategy, exploit the cultivation temperature, and at the same time explore the reward (food) through the IM strategy, with each change in temperature.

How do the worms acquire these two strategies? We found that in the starved condition, temperature and feeding were dissociated, and as a result the DM strategy disappeared, whereas the IM strategy was still applied (**Fig 4Ab**). According to these findings, we hypothesize that the DM strategy emerges as a consequence of associative learning between the cultivation temperature and food access; the IM strategy, however, could be innate. Further investigation of these hypotheses should be expected in the future.

### Comparison between WT and thermosensory neuron-deficient animals

In addition to WT worms, we identified the desirability functions of the thermosensory neuron (AWC and AFD)-deficient worms (**Fig 4B and C**). The AWC and AFD neurons are both known to sense the temporal derivative of temperature, *dT* [16,22,23]. However, the AWC-deficient worms showed a desirability function profile similar to that of WT worms (**Fig 4Bb**), whereas the AFD-deficient worms had a different profile (**Fig 4Cb**); the profile lacked the DM’s diagonal component and showed no bias along the *dT* axis. It can be assumed that an impaired AFD neuron prevents the worm from deciding whether an increase or decrease in temperature is favorable, which could lead to inefficient thermotactic migration. Thus, the AFD, but not the AWC neuron, is involved in oriented migration behavior based on temporal changes in temperature.

### Future perspective for neuroscience research

Finally, it is worth discussing future perspective of our IRL approach in neuroscience research focusing on higher-order animals beyond *C. elegans*. Over the last two decades, it has been clarified that dopaminergic neural activity in the ventral tegmental area (VTA) encodes the prediction error of reward [5], similar to temporal difference (TD) learning in RL [6]. Thus, an animal’s behavioral strategy should be associated with reward-based representation. It then has been widely believed that RL-like algorithm is processed within functionally networked cortical and subcortical areas, especially within basal ganglia [32–35] and amygdala [36,37], a group of brain nuclei heavily innervated by VTA dopaminergic neurons. Recent advances in neural recording technology enable us to monitor activities of neuronal population in freely-behaving animals, and the neuronal activities should be related to the reward-based representation of the strategy. However, there has been difficulty in knowing what are rewards for freely-behaving animals, especially internally-represented reward in the brain, rather than primitive rewards, e.g., foods. On the other hand, our IRL approach is able to identify the reward-based representation of the strategy, which allows analyses of neural correlates, such as comparing neural activities of freely-behaving animals with strategy-related variables calculated by IRL. Therefore, combination of neuroscience experiments and the IRL technology could contribute to discover neural substrate and its computational principle.

## Materials and Methods

### Reinforcement learning

Reinforcement learning (RL) is a machine learning model that describes how agents learn to obtain an optimal policy, that is, a behavioral strategy, in a given environment [4]. RL consists of several components: an agent, an environment, and a reward function. The agent learns and makes decisions, and the environment is defined by everything else. The agent continuously interacts with the environment, in which the state of the agent transits based on its action (behavior), and the agent gets a reward at the new state according to the reward function. The aim of the agent is to identify an optimal strategy (policy) that maximizes cumulative rewards in the long term.

In this study, the environment and the agent’s behavioral strategy were modeled as LMDP, one of the settings of RL [20]. An LMDP is characterized by the passive dynamics of the environment in the absence of control, and by controlled dynamics that reflect the behavioral strategy. Passive and controlled dynamics were defined by transition probabilities from state *s* to *s*’, *p*(*s*’|*s*) and *π*(*s*’|*s*), respectively. At each state, the agent not only acquires a reward, but also receives resistance to the passive dynamics (**Fig 1C**). Thus, the net reward is described as

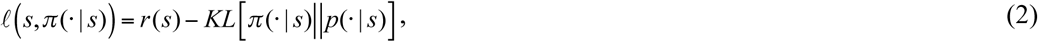

where *r*(*s*) denotes a state reward and *KL*[*π*(.|*s*)‖*p*(.|*s*)] indicates the Kullback–Leibler (KL) divergence between *π*(.|*s*) and *p*(.|*s*); this represents the resistance to the passive dynamics. The optimal policy that minimizes the cumulative net reward has been analytically obtained [20] as

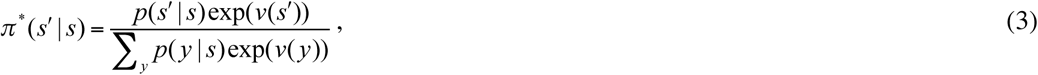

where asterisk means optimal, and *v*(*s*) is a value function, that is, the expected cumulative net rewards from state *s* toward the future:

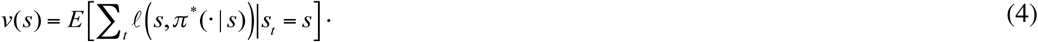

Here, we briefly show how to derive equation (3). First, the controlled dynamics was defined as

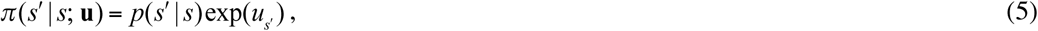

where the elements *u*_*s*_ of a vector **u** directly modulate the transition probability of passive dynamics. Note *π*(*s*’|*s*, **0**)= *p*(*s’*|*s*). Because of probability, equation (5) must satisfy

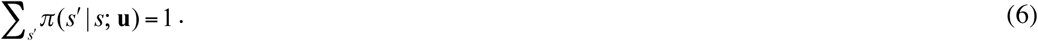

The value function can be rewritten by the Bellman equation:

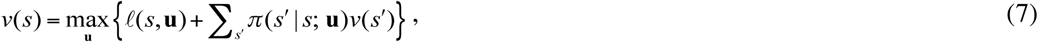

where *ℓ*(*s*,**u**) = *ℓ*(*s,π* (· | *s*; **u**)). The maximization in equation (7) subject to equation (6) by the method of Lagrange multipliers yields **u**^**\***^, which represents the optimal strategy. Substituting **u**^**\***^ in equation (5) becomes equation (3). In addition, substituting the optimal strategy (equation (3)) in the Bellman equation (7) and dropping the max operator lead to

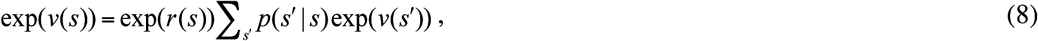

which satisfies Bellman’s self-consistency. Using this equation, the value function *v*(*s*) can be calculated from the reward function *r*(*s*), and vise versa. Please see [20] for full derivation.

### Inverse reinforcement learning (estimation of the value function)

To estimate the value function *v*(*s*), we assumed that the observed sequential state transitions {*s*_*t*_, *s*_*t*+1_}_*t*=1:*T*_ were generated by the stationary policy *π*^*\**^. We then maximized the likelihood of the sequential state transition:

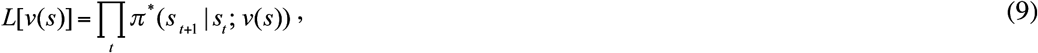

where *π*^*\**^(*s*_*t*+1_|*s*_*t*_; *v*(*s*)) corresponds to equation (3). This maximum likelihood estimation (MLE) was called OptV [21]. Based on the estimated value function, the primary reward function, *r*(*s*), can be calculated by using equation (8).

In our implementation, states were represented by tabular format, in which two-dimensional space (temperature and its temporal derivative) was divided as a mesh grid. Thus, our IRL requires a number of state trajectory data, which spans the entire mesh grid. In order to compensate for data limitation and noisy sensory systems, it is reasonable to assume that animals have value functions that are smooth in their state space. To obtain smooth value functions, we regularized MLE as

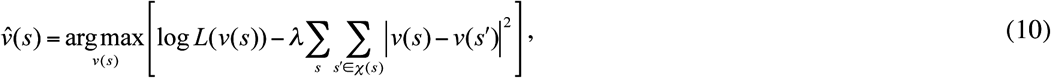

where the first term represents negative log-likelihood, and the second term represents a smoothness constraint introduced to the value function; a positive constant *λ* indicates the strength of the constraint, and *χ*(*s*) indicates a set of neighboring states of *s* in the state space. Notice that the cost function, the regularized negative log-likelihood, is convex with respect to *v*(*s*), which means there are no local minima in its optimization procedure.

### *Passive dynamics of thermotaxis in* C. elegans

To apply LMDP-based IRL to thermotactic behaviors of *C. elegans*, state *s* and passive dynamics *p*(*s*’|*s*) must be defined (phase 2 and 3 in **Fig 1D**). We previously found that the thermosensory AFD neuron encodes the temporal derivative of the environmental temperature [22], and thus assumed that the worm can sense not only absolute temperature *T*, but also the temporal derivative of temperature *dT/dt*. We therefore set a 2D state representation as (*T, dT*). Note that *dT/dt* is simply denoted as *dT*.

The passive dynamics were described by the transition probability of a state (*T, dT*) as

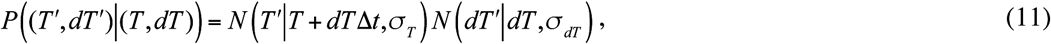

where 𝒩(*x*|*µ*, σ) indicates a Gaussian distribution of variable *x* with mean *µ* and variance σ, and *Δt* indicates the time interval of monitoring in behavioral experiments. This passive dynamics aspect can be loosely interpreted as the worms inertially migrating in a short time interval under a thermal gradient, but it may also be perturbed by white noise. Note that distribution for the passive dynamics can be arbitrary selected, and the choice of Gaussian was not due to mathematical necessity for the IRL.

### Artificial data

We confirmed that our regularized version of OptV (equation (6)) provided a good estimation of the value function using simulation data. First, we designed the value function of *T* and *dT* as the ground truth (**S2A Fig**), and passive dynamics through equation (7). Thus, the optimal policy was defined by equation (3). Second, we generated a time-series of state transitions according to the optimal policy, and separated these time series into training and test datasets. After that, we estimated the value function from the training dataset, varying the regularization parameter *λ* in equation (6) (**S2B Fig**). We then evaluated the squared error between the behavioral strategy based on the ground truth and the estimated value function, using the test dataset. Since the squared error on the test data was substantially reduced (by 88.1%) due to regularization, we deemed it effective for avoiding overfitting (**S2C Fig**).

### Cross-validation

In estimation of the value function, we performed cross-validation to determine *λ* in equation (10), and *σ*_*T*_ and *σ*_*dT*_ in equation (11), with which the prediction ability is maximized. We divided the behavioral time-series data equally into nine parts. We then independently performed estimation of the value function nine times; for each estimation, eight of the nine parts of the data were used for estimation, while the remaining part was used to evaluate the prediction ability of the estimated value function by the likelihood (equation (9)). We then optimized those parameters at which we obtained the lowest negative log-likelihood as averaged from the nine estimations.

### Surrogate method-based statistical testing

To check whether the DM and IM strategies were not obtained by chance, surrogate method-based statistical testing was performed under a null hypothesis that the worms randomly migrated under a thermal gradient with no behavioral strategy. We first constructed a set of artificial temperature time-series that could be observed under the null hypothesis. By using the iterated amplitude adjusted Fourier transform (IAAFT) method [38], we prepared 1000 surrogate datasets by shuffling observed temperature time-series (**S5A Fig**), while preserving the autocorrelation of the original time-series (**S5B Fig**). We then applied our IRL algorithm to this surrogate dataset to estimate the desirability functions (**S5C Fig**). To assess the significance of the DM and IM strategies, we calculated sums of the estimated desirability functions within the previously described horizontal and diagonal regions, respectively (**S5D Fig**). Empirical distributions of these test statistics for the surrogate datasets could then serve as null distributions (**S5E Fig**). For both DM and IM, the test statistic derived using the original desirability function was located above the empirical null distribution (*p*<0.001 for the DM strategy; *p*<0.001 for the IM strategy), indicating that both strategies were not obtained by chance but reflected an actual strategy of thermotaxis.

### C. elegans *preparation*

All worms were hermaphrodites and cultivated on OP50 as bacterial food using standard techniques [39]. The following strains were used: N2 wild-type Bristol strain, PY7505 *oyIs84[gcy-27p∷cz∷caspase-3(p17), gpa-4p∷caspase-3(p12)∷nz, gcy-27p∷GFP, unc-122p∷dsRed]*, IK2808 *njIs79[ceh-36p∷cz∷caspase-3(p17), ceh-36p∷caspase-3(p12)∷nz, ges-1p∷NLS∷GFP]* and IK2809 *njIs80[gcy-8p∷cz∷caspase-3(p17), gcy-8p∷caspase3(p12)∷nz, ges-1p∷NLS∷GFP]*. The ASI-ablated strain (PY7505) was a kind gift from Dr. Piali Sengupta [27]. The AFD-ablated strain (IK2809) and the AWC-ablated strain (IK2808) were generated by the expression of reconstituted caspases [40]. Plasmids carrying the reconstituted caspases were injected at 25 ng/µl with the injection marker pKDK66 (*ges-1p∷NLS∷GFP*) (50 ng/µl). Extrachromosomal arrays were integrated into the genome by gamma irradiation, and the resulting strains were outcrossed four times before analysis. To assess the efficiency of cell killing by the caspase transgenes, the integrated transgenes were crossed into integrated reporters that expressed GFPs in several neurons, including the neuron of interest, as follows: IK0673 *njIs2*[*nhr-38p∷GFP, AIYp∷GFP*] for AFD and IK2811 *njIs82*[*ceh-36p∷GFP, glr-3p∷GFP*] for AWC. Neuronal loss was confirmed by the disappearance of fluorescence; 100% of *njIs80* animals displayed the loss of AFD and 98.4% of *njIs79* animals displayed the loss of AWC.

### Thermotaxis assay

Thermotaxis (TTX) assays were performed as previously described [41]. Animals cultivated at 20 °C were placed on the center of an assay plate (14 cm × 10 cm, 1.45 cm height) containing 18 ml of TTX medium with 2% agar, and were allowed to freely move for 60 min. The center of the plate was adjusted to 17 °C, 20 °C, or 23 °C, to create three different gradient conditions, and the plates were then maintained at a linear thermal gradient of approximately 0.45 °C/cm.

### Behavioral recording

Worm behaviors were imaged using a CMOS sensor camera-link camera (8 bits, 4,096 × 3,072 pixels; CSC12M25BMP19-01B; Toshiba-Teli), a Line-Scan Lens (35 mm, f/2.8; YF3528; PENTAX), and a camera-link frame grabber (PCIe-1433; National Instruments). The camera was mounted at a distance above the assay plate that consistently produced an image with 33.2 µm per pixel. The frame rate of recordings was approximately 13.5 Hz. Images were captured and processed by a multi-worm Tracker [24], to detect worm bodies and measure behavioral parameters such as the position of the centroid.

## Acknowledgements

We thank Drs. Eiji Uchibe, Masataka Yamao, and Shin-ichi Maeda for their valuable comments. We are also grateful to Dr. Shigeyuki Oba for giving advice on statistical testing.

## Author Contributions

Conceptualization: HN SI

Data curation: SY MI

Formal analysis: SY HN

Funding acquisition: HN

Investigation: SY HN MI SN

Methodology: SY HN

Project administration: HN

Resources: SY MI SN

Software: SY

Supervision: HN

Validation: SY

Visualization: SY

Writing original draft: HN SY

Writing review & editing: HN SY YT MI SN IM SI

## Supporting figures

**S1 Fig:**
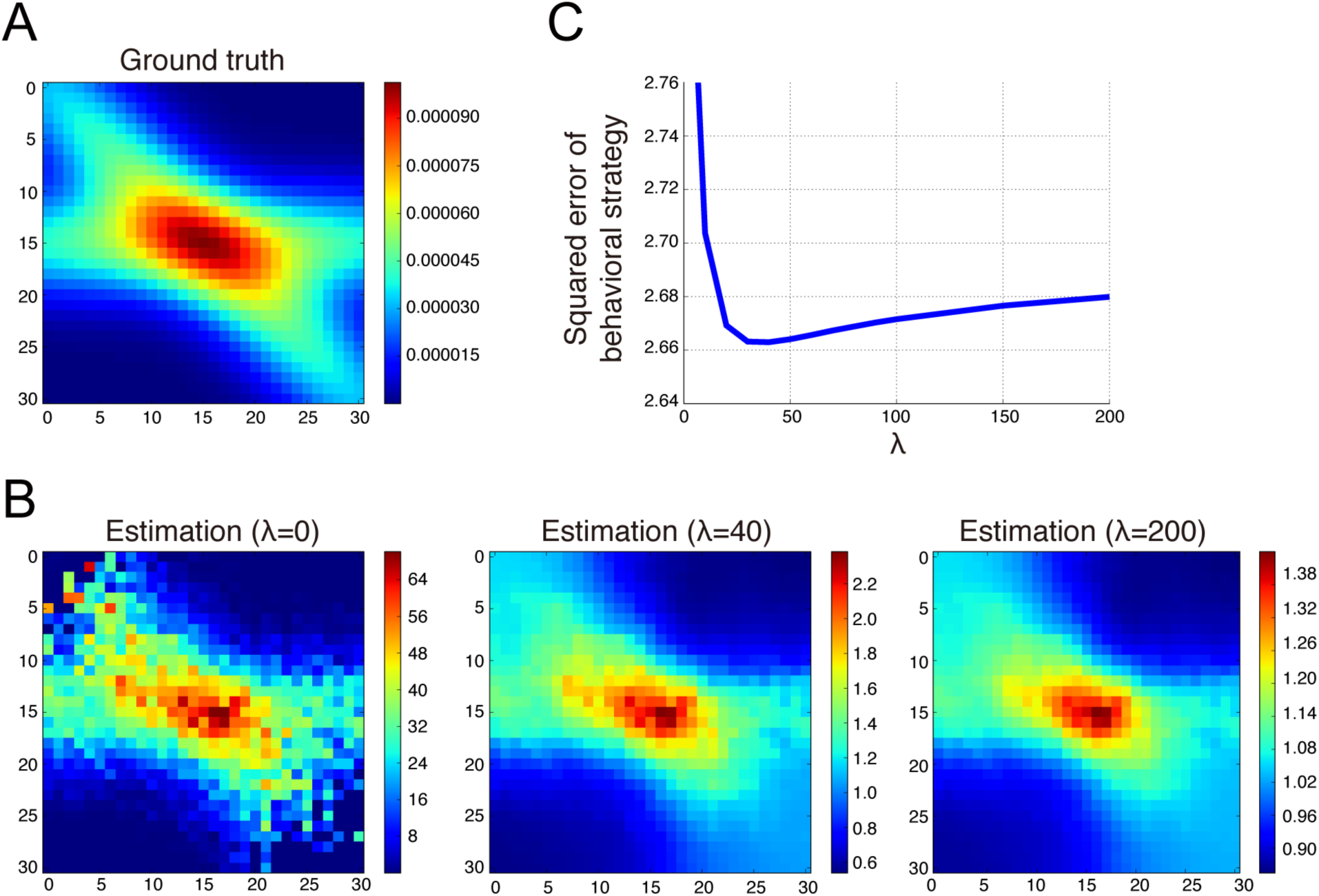
Validation of the regularized (OptV) estimation method in artificial data. **(A)** The desirability function corresponding to the ground truth value function used for generation of artificial data. Time-series data were artificially generated as training and test data sets by sampling equation (1), given the ground truth of the value function. **(B)** The desirability functions described by equation (6) under three different regularization parameters (*λ*) were visualized from the estimated value functions. **(C)** Squared error between the behavioral strategies based on the ground truth and estimated value functions using the test data set. The presence of an optimal *λ*, at which the minimal squared error is obtained, indicates that the regularization was effective for accurately estimating the value function.

**S2 Fig:**
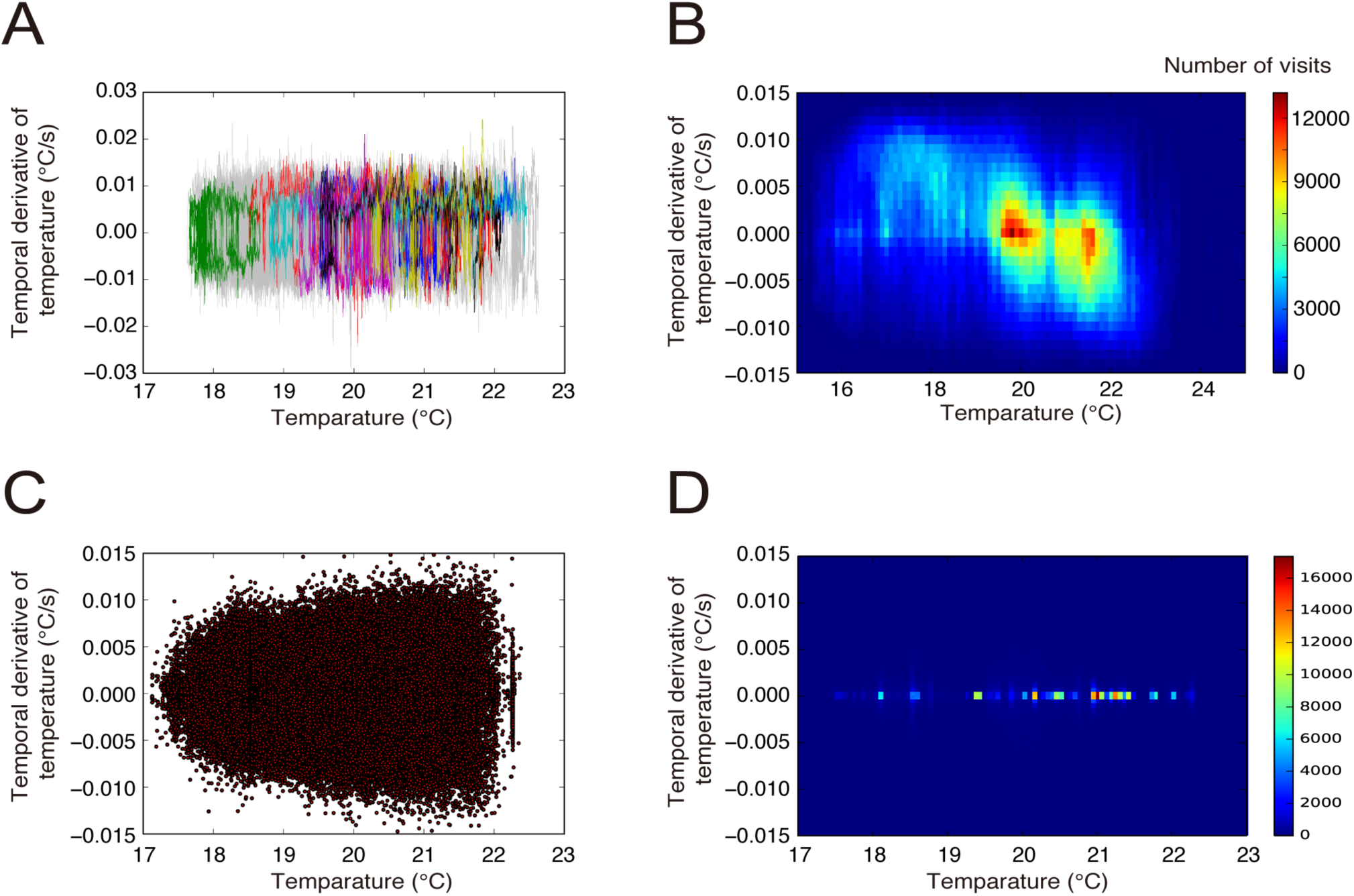
Behaviors in the *T*-*dT* space. **(A)** *T-dT* trajectories of fed WT worms. This is another representation of Fig. 2C and D. **(B)** Distributions of *T* and *dT* in all trajectories of fed WT worms. Notice that the distribution is substantially different from the desirability function (see Fig 3B). **(C)** Scatter plot of *T* and *dT* at 5 seconds before the moment of sharp turns. Correlation coefficient was 3.6e-10. Note that *dT* is 0 at the moment of sharp turns, because the worms stop for making large directional changes. **(D)** Histogram of the scatter plot in C.

**S3 Fig:**
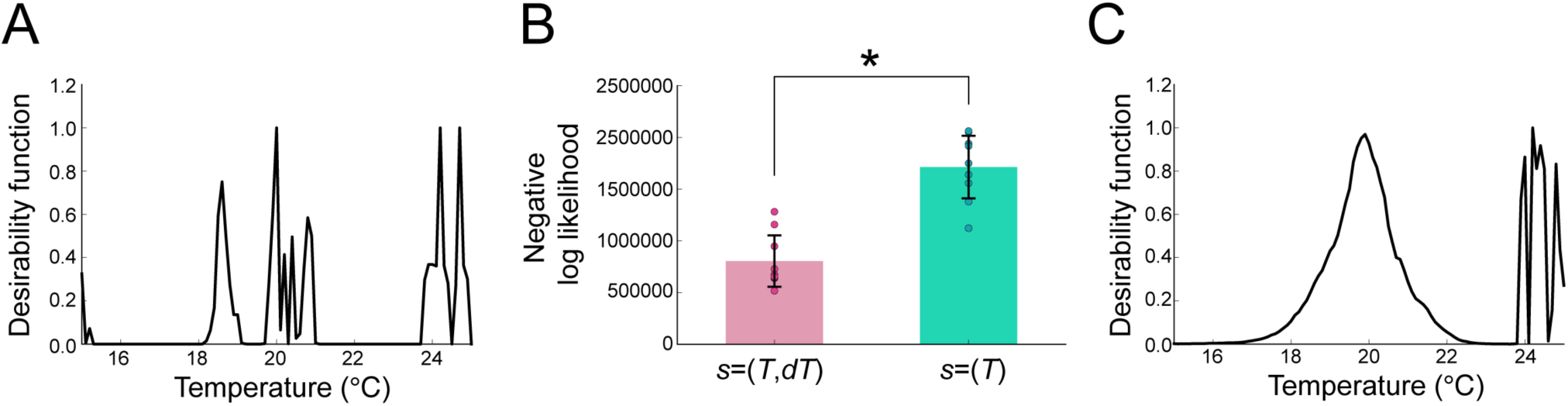
Inverse reinforcement learning (IRL) analysis with one-dimensional state representation. IRL was analyzed with one-dimensional state representation (*s*=(*T*)). **(A)** The desirability function was calculated using the estimated value function. In the estimation, the regularization parameter, *λ*, in equation (6) was optimized by cross-validation. **(B)** Prediction ability was compared between IRLs with *s*=(*T, dT*) and *s*=(*T*) using a cross-validation dataset. The negative log-likelihood of the behavioral strategies (equation (1)) with the estimated value function of both *T* and *dT* (see Fig 3B) was significantly smaller than that with the estimated value function of *T* alone (S3A Fig) (*p=*0.0002; Mann-Whitney U test). Thus, the behavioral strategy with *s*=(*T, dT*) was more appropriate than that with *s*=(*T*). **(C)** The desirability function became smoother as *λ* increased. This desirability function peaks around the cultivation temperature (20 °C).

**S4 Fig:**
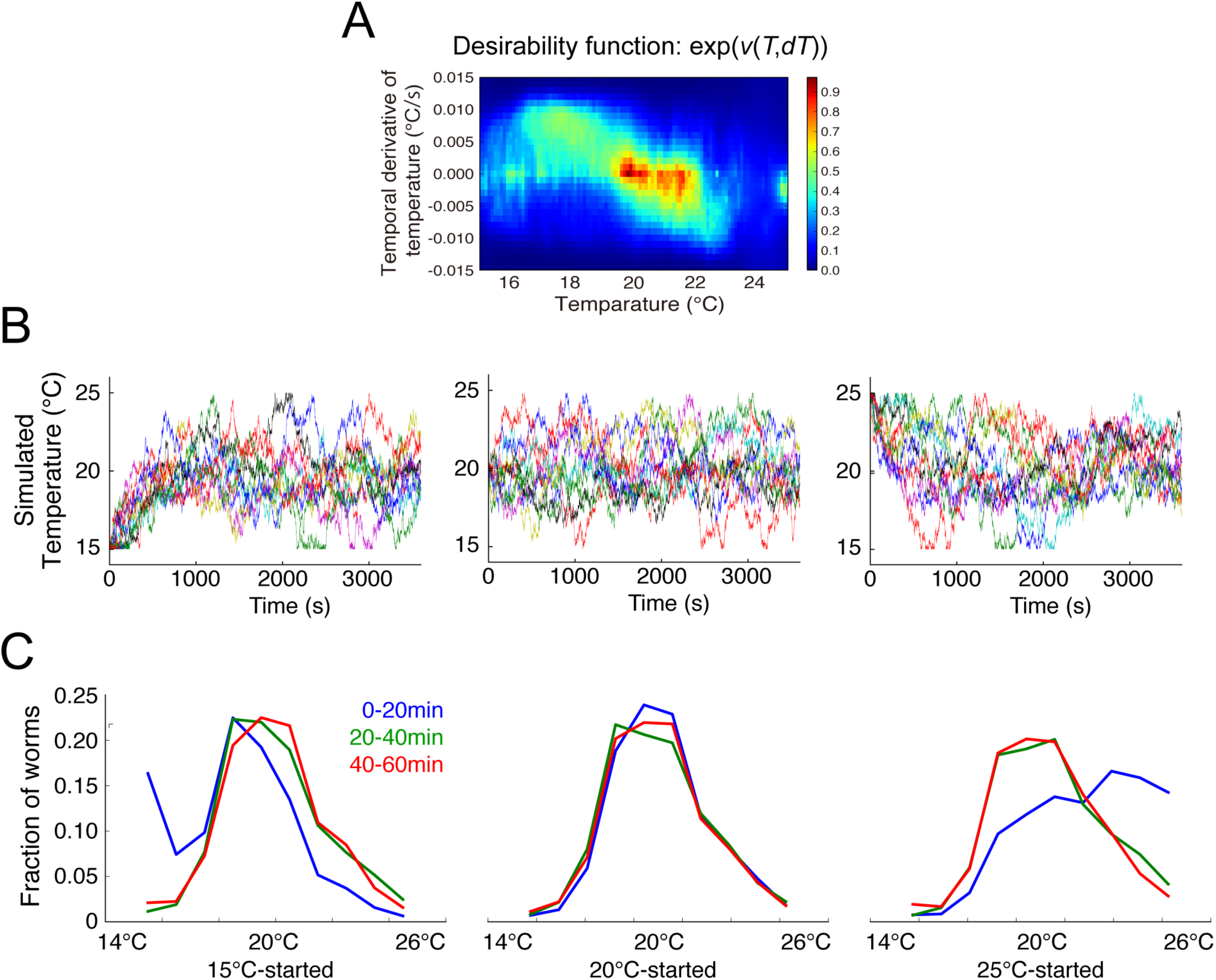
Reproduction of thermotaxis by simulating identified strategy. **(A)** The identified desirability function of the fed WT worms. This is identical to Fig. 3B. **(B)** Temperature time-series of simulated worms started from 15°C, 20°C or 25°C with 0°C/s. In the simulation, the state transition was sampled from equation (3) using the identified desirability function in (A). Different colored lines correspond to different simulation runs. **(C)** Temporal changes in distributions of 100 simulated worms. Notice that worm population converged around the cultivation temperature, i.e., 20°C.

**S5 Fig:**
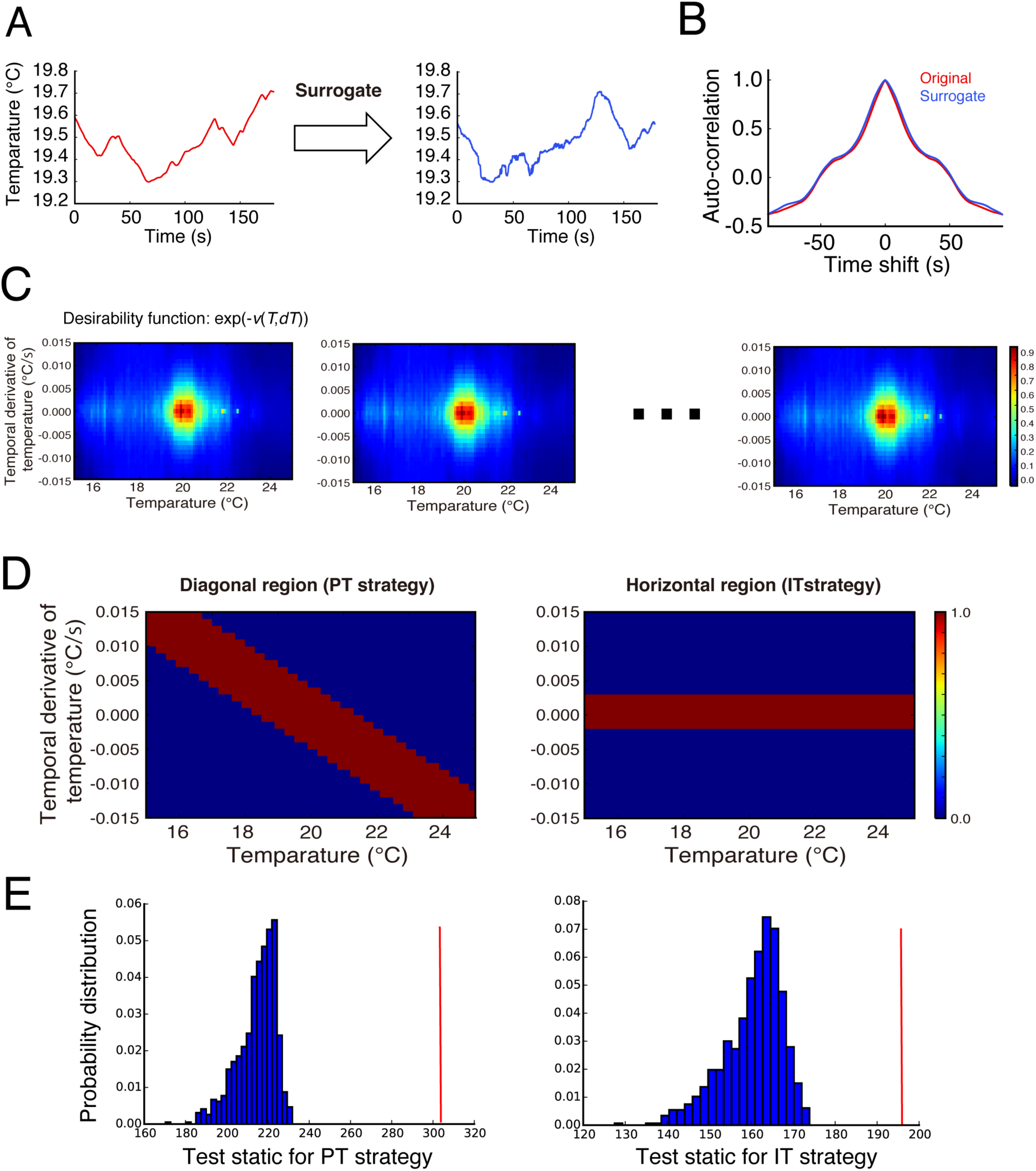
Statistical test for reliability of behavioral strategies with the surrogate method. The reliability of the directed migration (DM) and isothermal migration (IM) strategies (see Fig 3) was assessed by means of statistical testing with the null hypothesis that the worms randomly migrate with no behavioral strategy. **(A)** To generate time-series data under the null hypothesis, original time-series data of temperature (left panel) was surrogated by the IAAFT method (right panel). **(B)** Before and after the surrogation, the autocorrelations were almost preserved. **(C)** The desirability functions estimated from the surrogate datasets. **(D)** The DM and IM strategies correspond to the red-highlighted diagonal and horizontal regions of the desirability function, respectively. Within these regions, sums of the estimated desirability functions were calculated as test statistics. **(E)** Histograms of the empirical null distributions of the test statistics for the DM and IM strategies. The test statistics derived by the original desirability function (red arrows) are located above the empirical null distributions (*p*<0.001 for the PT strategy; *p*<0.001 for the IT strategy).

**S6 Fig:**
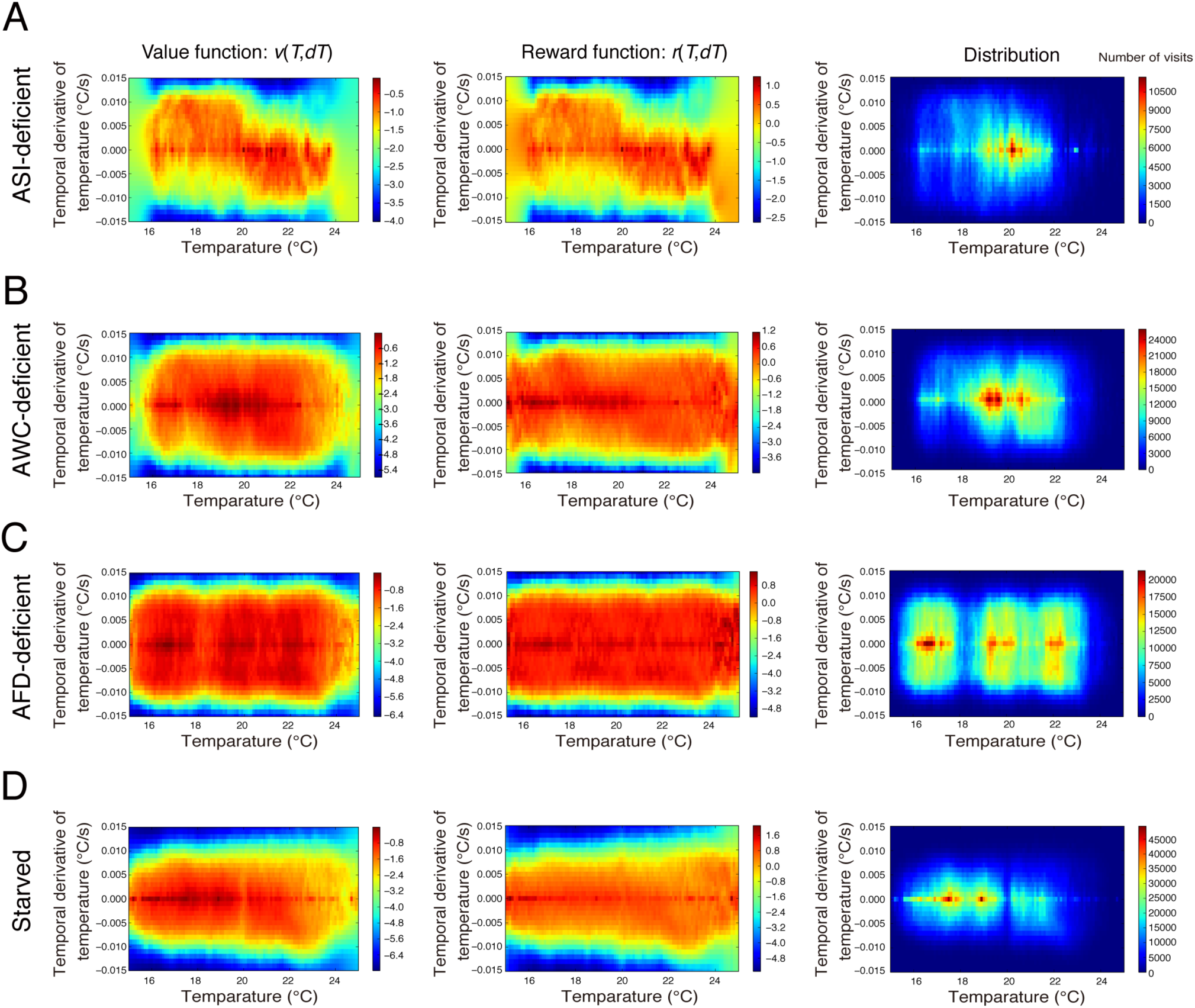
Estimated value/reward functions and state distributions. The estimated value functions, reward functions, and state distributions are depicted for the ASI-deficient worms **(A)**, the AWC-deficient worms **(B)**, the AFD-deficient worms **(C)** and starved WT worms **(D)**.

